# Prediction of the Human Pharmacokinetics of 30 Modern Antibiotics Using the ANDROMEDA Software

**DOI:** 10.1101/2023.03.28.534601

**Authors:** Urban Fagerholm, Sven Hellberg, Jonathan Alvarsson, Ola Spjuth

## Abstract

The ANDROMEDA software, based on machine learning, conformal prediction and a new physiologically-based pharmacokinetic model, was used to predict and characterize the human clinical pharmacokinetics of 30 selected modern small antibiotic compounds (investigational and marketed drugs). A majority of clinical pharmacokinetic data was missing. ANDROMEDA successfully filled this gap. Most antibiotics were predicted and measured to have limited permeability, good metabolic stability and multiple elimination pathways. According to predictions, most of the antibiotics are mainly eliminated renally and biliary and every other antibiotic is mainly eliminated via the renal route. Mean prediction errors for steady state volume of distribution, unbound fraction in plasma, renal and total clearance, oral clearance, fraction absorbed, fraction excreted renally, oral bioavailability and half-life were 1.3- to 2.3-fold. The overall median and maximum prediction errors were 1.5- and 4.8-fold, respectively, and 92 % of predictions had <3-fold error. Results are consistent with those obtained in previous validation studies and are better than with the best laboratory-based prediction methods, which validates ANDROMEDA for predictions of human clinical pharmacokinetics of modern antibiotic drugs, which to a great extent demonstrate pharmacokinetic characteristics challenging for laboratory methods (metabolic stability, limited permeability, efflux and multiple elimination pathways). Advantages with ANDROMEDA include that results are produced without the use of animals and cells and that predictions and decision-making can be done already at the design stage.

## Introduction

Antibiotic resistance is a serious emerging global health threat. Thus, there is a great demand to search for new effective and safe antibiotics.

Effective and safe dosing of new antibiotic drug candidates in humans requires that their pharmacokinetics (PK) is well predicted and defined. Prediction of PK for this class of compounds has generally been challenging, since many of them have limited passive permeability (P_e_) (with or without active uptake and efflux), are excreted to a significant extent via urine and bile and and/or are given at high dose levels (saturation PK).

Examples of antibiotics with limited P_e_ are amikacin, aztreonam, cefazolin, cefodizime, ceftriaxone, colistin, gentamicin, imipenem, kanamycin, neomycin and streptomycin, which all have maximum 5 % fraction absorbed (f_a_) and bioavailability (F) following oral dosing. Other antibiotics with limited passive P_e_, but that also utilize active intestinal uptake transport (via PEPT1), include β-lactam antibiotics ampicillin, cefexime and cefadroxil (their passive P_e_ correspond to <20 % f_a_, while their actual f_a_ is >50 %). Significant renal excretion occurs for, for example, fosfomycin, cefuroxime, cephradine, cefadroxil, cefcanel and cephalexin, which all have a fraction excreted in urine (f_e,renal_) of 90-100 % following intravenous dosing. Cefoperazone, ceftriaxone, erythromycin, mezlocillin and piperacillin are among antibiotics that have a fraction excreted in bile (f_e,bile_) of at least 20 % following intravenous dosing. The highly lipophilic clofazimine and bedaquiline (both used to treat tuberculosis) bind extensively to fat tissue and plasma proteins and have a half-life (t_½_) of several months.

Poor f_a_ and F prevent some antibiotic drugs to be suitable for oral dosing, and poor uptake and significant elimination via urine and feces means that a high fraction of dose (and sometimes also comparably large amount) is transported out into the environment. Thus, an ambition to enhance the oral uptake and reduce the fractional excretion of antibiotics is beneficial for the environment. Another implication of poor oral absorption and major excretion via kidneys and bile is limited applicability of *in vitro* PK screening systems such as Caco-2 (uncertain f_a_- and F-predictions) and hepatocytes (metabolism rate below the limit of quantification). This was shown for ciprofloxacin (considerable underprediction of f_a_ from Caco-2 P_e_) and piperacillin (metabolic stability not possible to quantify with human microsomes) (Fagerholm and Lundqvist, unpublished data).

ANDROMEDA by Prosilico is a multi-validated integrated *in silico*-based prediction software for human clinical PK.^1-4^ It is based on conformal prediction (CP), which is a methodology that sits on top of machine learning methods and produce valid levels of confidence,^5^ and a novel physiologically-based pharmacokinetic (PBPK) model.^1^ See Alvarsson et al. (2021) for a more extensive introduction to CP.^6^ The software has been applied for prediction of human clinical PK, including fraction absorbed f_a_, F, t_½_, *in vivo* dissolution potential, unbound fraction in plasma (f_u_), intrinsic metabolic clearance (CL_int_), CL, oral CL (CL/F), steady-state volume of distribution (V_ss_), f_e,renal_ and f_e,bile_.^2-4^ In a major benchmarking study it outperformed laboratory methods in predictive accuracy and range.^3^ ANDROMEDA predictions have also been approved by the German authority BfArM for use and main preclinical PK-source in clinical trial applications.

In 2017, 48 new antibiotic compounds with the potential to treat serious bacterial infections were in clinical development.^7^ In an article by Al-Tawfiq et al., 20 antibiotics in the pipeline (phase III) or new on the market 2017-2020 were reviewed.^8^ Compounds from these selections opened up for the opportunity to further validate ANDROMEDA and to characterize the PK of modern antibiotic compounds. The main aim of this study was to validate ANDROMEDA for prediction and characterization of the human clinical PK (with main focus on elimination routes) of modern antibiotics.

## Methods

### ANDROMEDA PK-Prediction Software

For a description of ANDROMEDA (including underlying CP models, algorithms, molecular descriptors, parameters and performance) see Fagerholm et al. 2022 and 2023.^1-4^ The software predicts 30 human PK-parameters, including V_ss_, f_u_, CL, CL_int_, f_e,renal_, f_e,bile_ (considering enterohepatic circulation), f_a_, F, t_½_ and CYP substrate (2C9, 2D6 and 3A4) and efflux transporter (MDR-1, BCRP and MRP2) specificity.

ANDROMEDA is mainly applicable for compounds with MW 100 to 700 g/mol and for non-saturated conditions (not a high doses). There are groups of compounds for which the *in silico* models do not work, including metals and quaternary amines, and have limited use, for example, hydrolysis sensitive compounds and drugs binding covalently and/or to DNA.

None of the new antibiotics selected for the study was included in the training sets of the used CP models. Thus, every prediction was a forward-looking prediction where each compound was unknown to the models.

Where possible for marketed antibiotics, prediction accuracy/errors were estimated as predicted/observed or observed/predicted values (ratios≥1 were selected). For investigational drugs, only qualitative (yes/no) data for renal and biliary excretion are presented.

### Compound Selection and Clinical PK-Data

30 small investigational antibiotics (n=18) and antibiotics marketed 2017-2020 (n=12) were explored and selected for the investigation (Table 1). The minimum, median, mean and maximum molecular weights for this selection were 277, 405, 461 and 937 g/mol, respectively. Antibiotics belonged to various classes: Fluoroquinolones, β-lactams, β-lactamase inhibitors, aminoglycosides, tetracyclines, pleuromutilins, cephalosporins, nitroimidazooxazines, DHFR inhibitors, 4th generation macrolides, triazaacenaphthylenes, spiropyrimidinetriones and oxazolidinones. PK-data of marketed antibiotics were taken from FDA-label documents (Clinical Pharmacology sections).

**Table 1.**
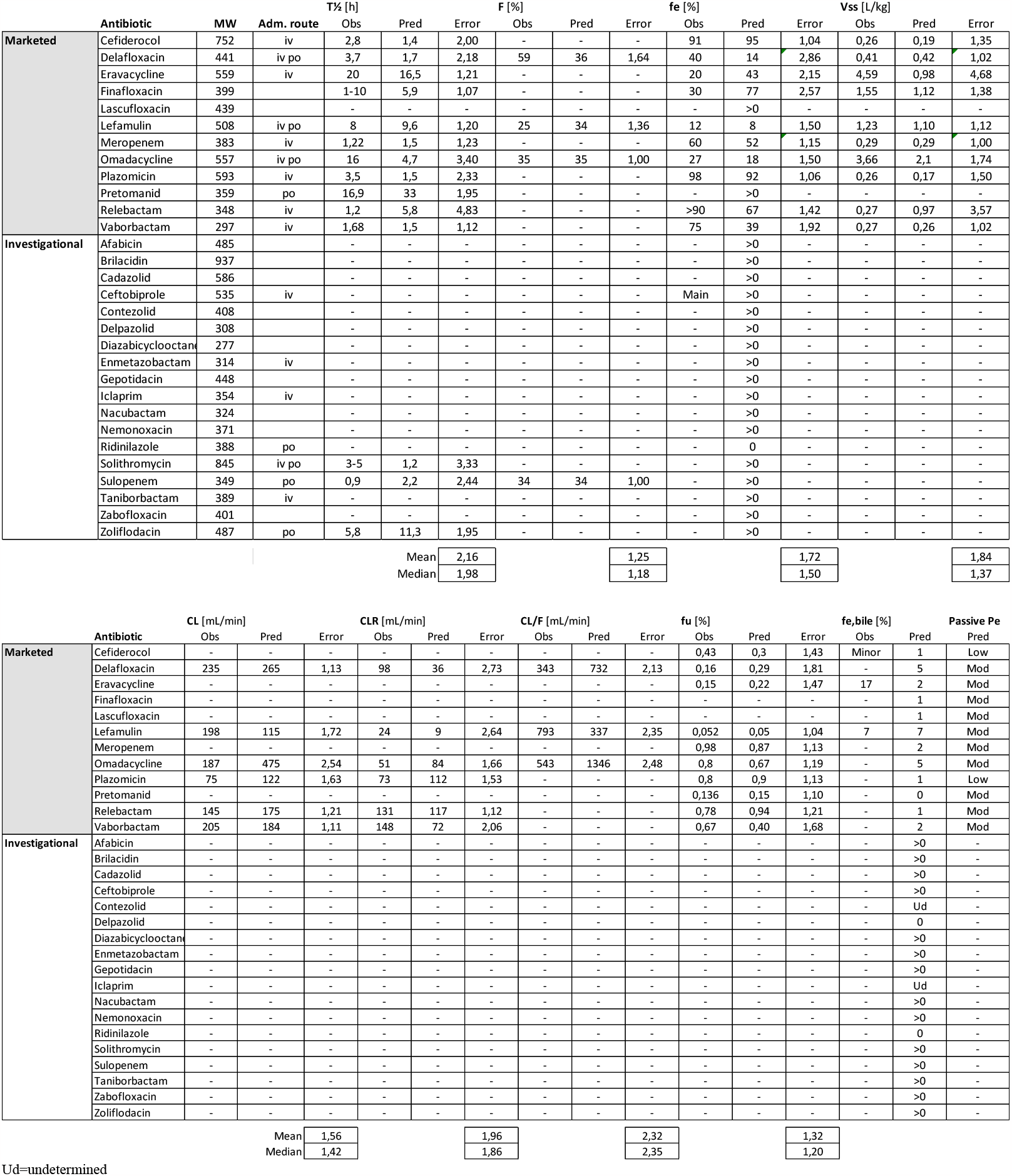
30 small investigational antibiotics and antibiotics marketed 2017-2020 and their observed and predicted human clinical pharmacokinetics.

## Results and Discussion

The amount of clinical PK-data for the antibiotics was limited – V_ss_ (n=10; range 0.3-4.6 L/kg; median 0.3 L/kg), f_u_ (n=10; 0.05-0.98; median 0.55), f_a_ (n=0), F (n=4; 0.25-0.59), CL (n=6; 75-235 mL/min), CL_R_ (n=6; 24-148 mL/min), CL/F (n=3; 343-793 mL/min), t_½_ (n=14; 0.9-20 h; median 3.6 h), f_e,bile_ (n=0; rough estimates were found for 3 antibiotics), f_e,renal_ (n=11; range 0.12-0.98; median 0.57).

For modern antibiotics in general, V_ss_, CL_int_ and t_½_ are low/short and f_u_, f_e,renal_ and f_e,bile_ high compared to other types of modern and traditional drugs.^3^ For 4 antibiotics, the t_½_ was less than 2 h. Rapid elimination from the body implies needs to administer at higher doses, with shorter time intervals and/or as continuous infusion.

ANDROMEDA was capable of predicting the human clinical PK for all the antibiotics. Prediction results are shown in Table 1. Median prediction errors for V_ss_, f_u_, CL, CL_R_, CL/F, F, f_e,renal_ and t_½_ were 1.4-, 1.2-, 1.4-, 1.9-, 2.3-, 1.2-, 1.5- (9 % absolute error) and 2.0-fold, respectively. Corresponding mean errors were 1.8-, 1.3-, 1.6-, 2.0-, 2.4-, 1.2-, 1.7- (18 % absolute error) and 2.2-fold, respectively. Maximum errors were 4.7-, 1.8-, 2.5-, 2.7-, 2.5-, 1.6-, 2.9- (47 % absolute error) and 4.8-fold, respectively.

For eravacycline, lefamulin and cefiderocol, a minor fraction (up to 17 %) unchanged substance was found in faeces following intravenous dosing. This was consistent with up to 7 % predicted f_e,bile_ for these antibiotics.

The results for CL and CL/F indicate that the dose and exposure of antibiotic candidate drugs can be well predicted using the software (with on average ca 2-fold prediction error).

Overall prediction errors (median 1.5-fold, mean 1.8-fold, max 4.8-fold, 71 % with <2-fold error; 92 % with <3-fold error; n=63) are in line with previous results, which further validates ANDROMEDA. The results are equally good as or better than those obtained with best of comparable laboratory methods.^3,9,10^

According to predictions for antibiotics, most of them are mainly (≥50%) eliminated renally and biliary, 47 % are mainly (≥50%) eliminated via the renal route, 97 % are excreted renally, 83 % are excreted via bile (in an internal validation the model for bile excretion gave correct binary predictions in 97 % of cases), 93 % have low/moderate passive P_e_ (only one has very high passive P_e_), 43 % are effluxed by MDR-1, BCRP and/or MRP2, 20 % have limited gastrointestinal dissolution potential, 70-83 % have a CL_int_ that is normally problematic to quantify with human hepatocytes (approximately <500-1000 mL/min) and 60 % are not metabolised by CYPs 2C9, 2D6 and/or 3A4 (Figure 1).

**Figure 1.**
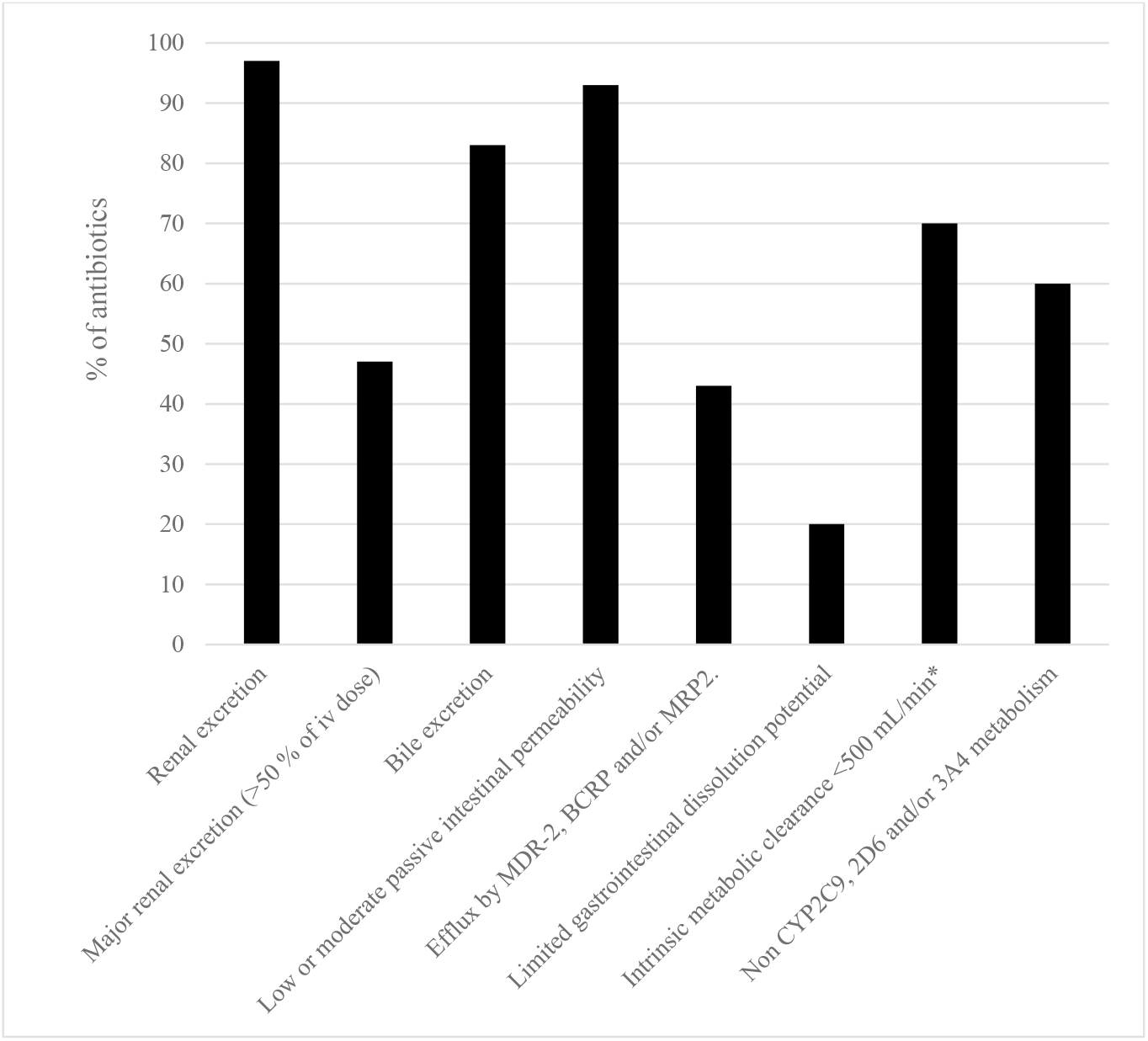
The percentage of modern antibiotics with certain predicted pharmacokinetics characteristics. *below the general limit of quantification for the conventional human hepatocyte assay.

Significant renal and biliary clearance and limited metabolic rate (often out of reach for the conventional hepatocyte assay) and P_e_ (challenging for Caco-2) demonstrates the requirement of methods to predict these properties for antibiotics *in vivo* in man. ANDROMEDA has this capacity (our *in silico* methodology also enables prediction of f_a_ for actively absorbed compounds^3^). It also shows a general trend that antibiotics to a major extent leaves the body in unchanged form out into the environment. Only one of the antibiotics does not undergo elimination via kidneys and bile (according to predictions).

In conclusion, the results are consistent with those obtained in previous validation studies and validates ANDROMEDA for predictions of human clinical PK of modern antibiotic drugs, which to a great extent demonstrate PK-characteristics challenging for laboratory methods (metabolic stability, limited P_e_, efflux and multiple elimination pathways). Advantages with ANDROMEDA include that PK-results are produced without the use of animals and cells and that predictions and decision-making can be done already at the design stage.

